# Ubiquitin isopeptidase and esterase activities have distinct catalytic requirements

**DOI:** 10.64898/2026.10.20.760496

**Authors:** Rachel E Lacoursiere, Paul P Geurink, Jonathan N Pruneda

## Abstract

Post-translational ubiquitin signaling is reversed by dedicated proteases termed deubiquitinases, many of which utilize a cysteine-based catalytic triad to cleave the isopeptide linkage. This activity represents canonical isopeptidase activity that has been mechanistically well characterized. By contrast, non-canonical ubiquitination, linked instead by an ester bond, is reversed by esterase activity through a relatively less well understood process. Several deubiquitinases, including examples from pathogenic bacteria and humans, have been shown to harbor dual isopeptidase and esterase activities. Here, we show that while both functions can take place within the same active site, they exhibit distinct catalytic requirements. In fact, the catalytic cysteine itself is not strictly required for ubiquitin esterase activity, as highlighted by substantial residual activities in both bacterial and human deubiquitinases following the typical Cys-to-Ala substitution. A Cys-to-Thr substitution, meanwhile, effectively eliminates both isopeptidase and esterase activity. Further illustrating distinct catalytic requirements, cryptic ubiquitin esterase activities were identified in OTUD6B and OTULINL, two reportedly inactive human (pseudo-) deubiquitinases. Overall, our work establishes new perspectives and approaches for studying the regulation of non-canonical ester-linked ubiquitination.

## Introduction

Canonically, ubiquitin (Ub) acts as a post-translational modifier that targets one or more lysine (Lys, K) residues within a proteinaceous substrate (Swatek & Komander, 2016). These Lys linkages are formed through stable isopeptide bonds between the C-terminal carboxylate of Ub and the amine side chain of the substrate Lys residue. Alternative linkages formed through non-canonical ubiquitination, loosely defined as modification of non-Lys residues, have also been described and recently expanded to even include non-proteinaceous targets (Kelsall, 2022; Squair & Virdee, 2022). Non-canonical ester-linked ubiquitination has been observed on serine/threonine (Ser/Thr, S/T) residues within proteins, as well as on hydroxyl groups of non-proteinaceous targets including lipids, sugars, and nucleic acids. Ester-linked substrates are difficult to identify and study due to the inherent lability of the ester bond as well as the lack of enzymes known to regulate these signals. While the cellular outcomes are still emerging, ester-linked ubiquitination has been linked to processes such as xenophagy, glycogen metabolism, and ER-associated degradation (ERAD) (Jochem *et al*, 2026; Otten *et al*, 2021; Wang *et al*, 2007; Wang *et al*, 2009; Yip *et al*, 2026).

Nearly 1000 human writers, readers, and erasers are known to regulate ubiquitin signaling, but only a small fraction have been described to function in non-canonical ubiquitination. Several writers including the E2 Ub-conjugating enzyme UBE2J2 and E3 Ub ligases MYCBP2, ARIH1, and HOIL1 have been shown to ubiquitinate Ser/Thr residues in protein substrates (Kelsall *et al*, 2022; Kelsall *et al*, 2019; McCrory *et al*, 2022; Pao *et al*, 2018; Purser *et al*, 2023; Wang *et al*., 2007). Additionally, the E3 ligases HOIL1, ARIH1, and RNF213 can catalyze ester-linked ubiquitination of non-proteinaceous substrates including carbohydrates, small molecules, and bacterial lipids (Jochem *et al*., 2026; Kelsall *et al*., 2022; Otten *et al*., 2021; Wang *et al*, 2025; Yip *et al*., 2026; Yoshida *et al*, 2024). The Deltex family of E3 ligases target the small molecule NAD+ and other ADP-ribose containing macromolecules for ubiquitination (Ahmed *et al*, 2020; Chatrin *et al*, 2020; Dearlove *et al*, 2024; Zhu *et al*, 2024; Zhu *et al*, 2022). Deltex E3 enzymes can modify protein-conjugated ADP-ribose as well, creating a hybrid mono-ADP-ribose-ubiquitin ester. Two proteins, RNF114 and RNF166, act as readers of this hybrid ubiquitin ester by non-covalently binding and subsequently elongating a Lys11-linked ubiquitin chain (Kloet *et al*, 2025; Lacoursiere *et al*, 2025; Perrard *et al*, 2025). Finally, several deubiquitinases (DUBs) can reverse this chemistry *in vitro*, including several members of the Ub-specific protease (USP) family. Interestingly, the Josephin-containing proteins JOSD1 and JOSD2 display innate specificity *in vitro* toward a Ub-Thr substrate compared to several Lys-linked Ub substrates (De Cesare *et al*, 2021; Ritorto *et al*, 2014). The biological relevance of these Ub esterase activities in cells remains to be determined.

Although there are multiple classes of DUBs, each with distinct protein folds and active-site conformations, all except the metalloproteases are cysteine proteases that utilize a catalytic triad comprised of a cysteine (Cys, C) nucleophile, a histidine (His, H) base, and a polar residue such as aspartate (Asp, D), asparagine (Asn, N), or in some cases a serine (Ser, S) (Nijman *et al*, 2005). Canonically, the Cys side chain thiol is deprotonated by the His side chain imidazole ring, which is in turn stabilized in its protonated state by the nearby polar Asp/Asn residue. Nucleophilic attack of a ubiquitinated substrate by the catalytic Cys yields a tetrahedral oxyanion intermediate that is stabilized by what is termed the oxyanion hole, which in some DUB classes contains an Asn or glutamine (Gln, Q) residue N-terminal to the catalytic Cys. Next, collapse of the tetrahedral intermediate liberates the C-terminus of ubiquitin from the substrate Lys, which can then be protonated and diffuse away, leaving behind a DUB∼Ub acyl intermediate. Finally, to regenerate the active site, a water molecule activated by the catalytic triad His hydrolyzes the C-terminus of ubiquitin from the catalytic Cys through a second tetrahedral intermediate (Mevissen & Komander, 2017). To study the biochemical, cellular, and physiological function of DUBs, common practice is to substitute the catalytic Cys residue for an alanine (Ala, A) or the more structurally analogous Ser, which ablates the nucleophile that initiates the deubiquitination reaction. This approach has proven useful for studying the structure, mechanism, and function of DUBs including OTULIN, OTUD2, MINDY1, MINDY2, CYLD, USP30, Cezanne, and others (Abdul Rehman *et al*, 2021; Abdul Rehman *et al*, 2016; Gersch *et al*, 2017; Keusekotten *et al*, 2013; Mevissen *et al*, 2013; Mevissen *et al*, 2016; Sato *et al*, 2015). In some cases, an arginine (Arg) substitution has been incorporated instead, to ablate nucleophilic function and simultaneously sterically restrict ubiquitin binding (Morrow *et al*, 2018).

Interestingly, several DUB family members show little or no activity toward canonical isopeptide-linked Ub substrates, including ALG13, OTUD6B, and OTULINL (FAM105A) (Ceccarelli *et al*, 2019; Mevissen *et al*., 2013; Ritorto *et al*., 2014). While all three of these examples structurally resemble a DUB, OTULINL uniquely lacks conservation of a catalytic Cys and is thus classified as a pseudo-DUB. Instead, OTULINL harbors an Asp at the site analogous to Cys129 in OTULIN (Keusekotten *et al*., 2013; Mevissen *et al*., 2013). Interestingly, members of the Machado-Joseph disease DUBs (MJDs), including JOSD1, were also characterized as weak or inactive enzymes until a comprehensive study of human DUBs against Ub-Lys and Ub-Thr substrates was conducted (De Cesare *et al*., 2021; Ritorto *et al*., 2014). This study demonstrated that while some DUB families exhibit dual isopeptidase and esterase activity, MJD family members JOSD1, JOSD2, and ATXN3 displayed dramatically higher esterase activity toward the Ub-Thr substrate, highlighting the importance of testing several types of ubiquitinated substrates. While ALG13 and OTULINL were not included in this new screen, OTUD6B was reportedly inactive against the tested substrates (De Cesare *et al*., 2021).

We previously identified TssM, a DUB secreted by *Burkholderia pseudomallei* during infection that protects the bacterium from RNF213-mediated ubiquitination and subsequent xenophagy (Szczesna *et al*, 2024). We showed that TssM possesses both canonical isopeptidase and non-canonical esterase DUB activities, and through point mutation created an esterase-specific version of the enzyme that has served as a useful tool in the discovery of additional ester-linked ubiquitin substrates (Bejan *et al*, 2025; Douglas *et al*, 2026). Here, we show that the dual catalytic activities of TssM can utilize distinct mechanisms of deubiquitination. We highlight that individual components of the catalytic triad required for canonical isopeptidase activity are dispensable for the esterase activity of TssM. Remarkably, we find that the classical Ala mutation of the catalytic Cys residue only abrogates isopeptidase activity of TssM, while a threonine (Thr, T) mutation is required to eliminate both isopeptidase and esterase function. With these insights, we revisit a series of human DUBs that were reportedly inactive and/or pseudo-enzymes and identify cryptic Ub esterase activities in OTUD6B and OTULINL. Finally, we propose that esterase activity in DUBs can arise from direct water-mediated hydrolysis within the active site, without formation of a DUB∼Ub covalent intermediate, and that either combinatorial or a catalytic Cys-to-Thr substitution is required to eliminate esterase activity.

## Results

### An alanine substitution at C292 in TssM retains activity toward ester-linked ubiquitin substrates

As part of our examination of *Burkholderia* TssM, we introduced an Ala substitution at the catalytic Cys residue (C292), hereafter referred to as CA, which markedly reduced DUB activity at the low enzyme concentrations used for comparison to the wildtype protein (Szczesna *et al*., 2024). At elevated enzyme concentrations (0.5-5 µM), we observed increased fluorescence polarization indicative of a noncovalent TssM:Ub interaction, but no capacity to cleave a canonical isopeptide-linked Ub-Lys substrate (Figure 1A). Under the same conditions, we were surprised to observe that TssM CA retained some Ub esterase activity toward the Ub-Ser and Ub-Thr substrates in a concentration-dependent manner (Figure 1B, C). This activity persisted across multiple rounds of sequence validation and protein purification, eliminating the possibility of any contaminating wildtype enzyme. When incubated with the Ub-propargylamide (Ub-PA) probe that covalently captures the catalytic Cys of DUBs (Ekkebus *et al*, 2013), TssM CA failed to form a covalent Ub adduct over the course of 20 hours (Figure 1D), consistent with a loss of the reactive Cys nucleophile. Compared to the catalytic efficiency of wildtype TssM we derived previously, the CA version of TssM exhibited reduced but still appreciable k_cat_/K_M_ for the Ub-Ser and Ub-Thr substrates (Figure 1E). Therefore, while a CA substitution may abolish canonical isopeptidase activity of a dual activity DUB such as TssM, esterase activity can be retained, underlining a crucial difference in the catalytic hydrolysis of Ub isopeptides and Ub esters.

**Figure 1.**
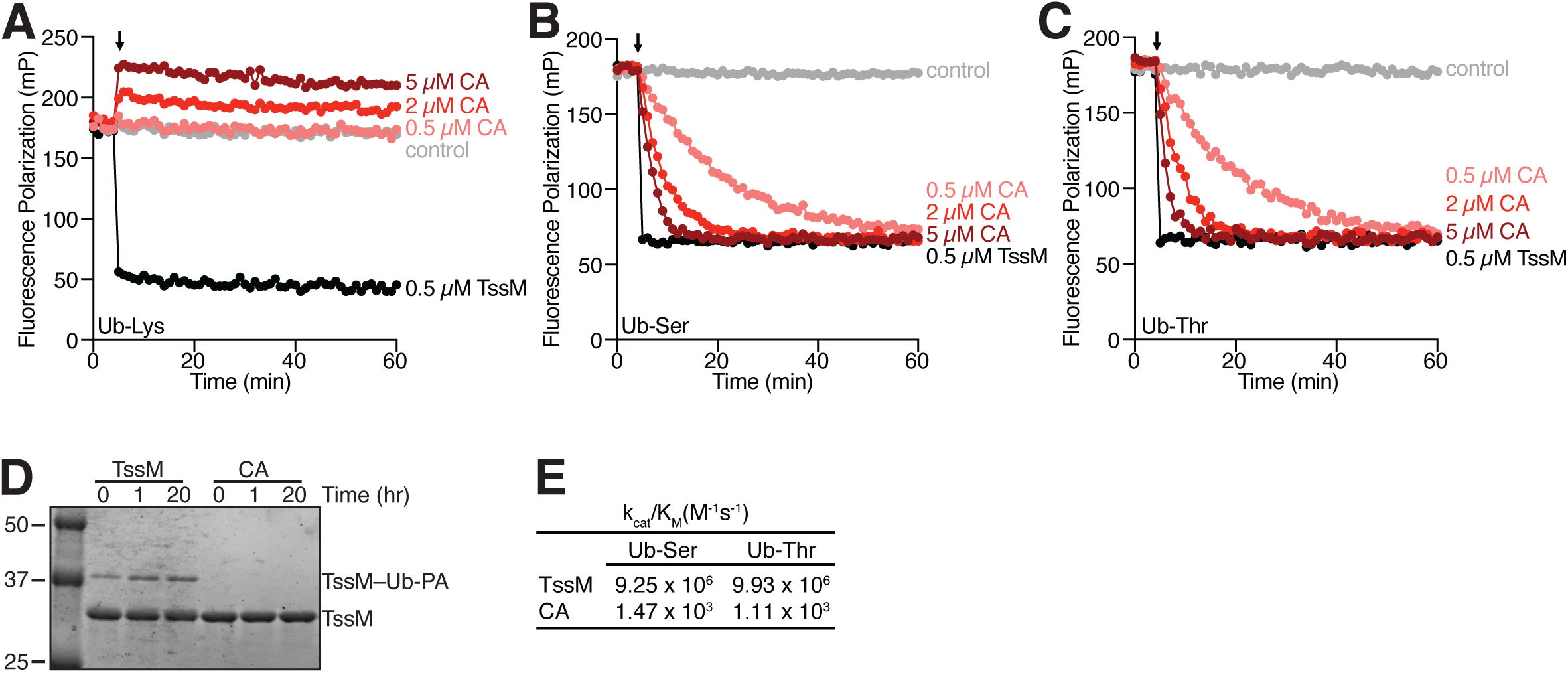
The standard Cys-to-Ala substitution at the TssM active site does not ablate its esterase activity. Fluorescence polarization experiments measuring activity of the CA version of TssM against the **(A)** Ub-Lys, **(B)** Ub-Ser, and **(C)** Ub-Thr substrates at several enzyme concentrations. The timepoint where the DUB was added to the reaction is marked by an arrow in all panels. **(D)** Time course showing the covalent modification of TssM and the CA version of TssM with the Ub-PA activity-based probe. **(E)** Catalytic efficiency of the TssM CA mutant against the Ub-Ser and Ub-Thr substrates. Values for wildtype TssM, reported previously (Szczesna *et al*., 2024), are shown for comparison.

### The ubiquitin esterase activity of TssM does not require the canonical catalytic triad

Canonically, Cys-based DUBs rely upon a properly aligned Cys/His/X (where X is a polar residue) catalytic triad for activity (Nijman *et al*., 2005). The catalytic triad of TssM is aligned in crystal structures bound to ubiquitin and consists of Cys292, His426, and Asp441 (Hermanns *et al*, 2024; Szczesna *et al*., 2024). The oxyanion hole, poised to stabilize the tetrahedral intermediate state, is formed by Asn286. To probe the requirement of these additional catalytic residues, we evaluated the consequences of Ala substitutions using a Ub-PA reactivity experiment. As expected, all versions of TssM tested that retained Cys292 were capable of covalent modification with Ub-PA, whereas TssM proteins containing a CA substitution were incapable of Ub-PA modification (Figure 2A).

**Figure 2.**
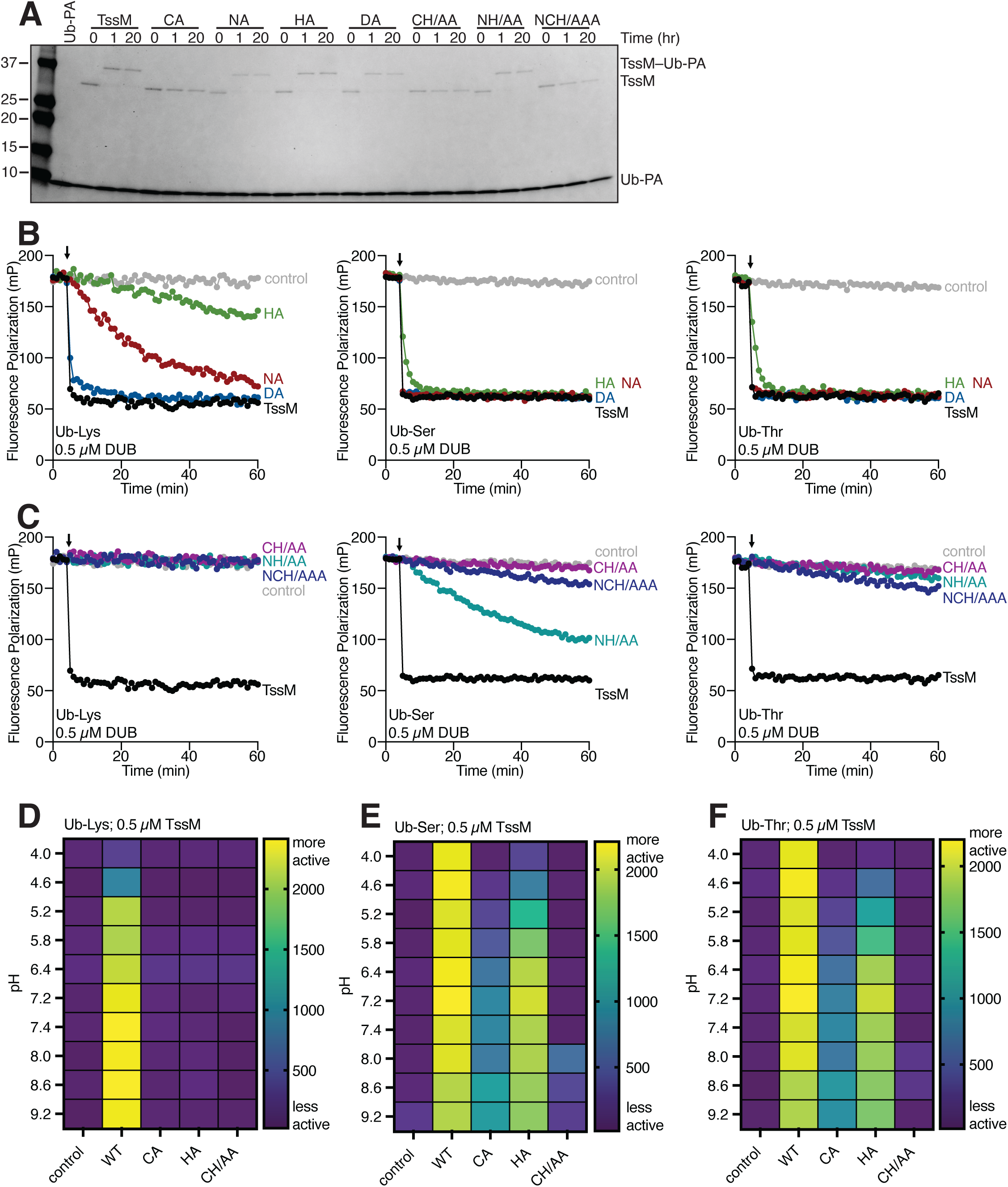
Cys292 or His426, individually, are sufficient to promote esterase activity in TssM. **(A)** Time course showing Ub-PA modification of the indicated versions of TssM. Fluorescence polarization experiments were used to show the activity of each variant of TssM against the Ub-Lys, Ub-Ser, and Ub-Thr substrates for **(B)** single alanine substitution of catalytic residues (His426, HA; Asp441, DA) and the oxyanion hole (Asn286, NA), and **(C)** double or triple alanine substitution for several catalytic residues. Addition of the DUB is marked by the black arrow for all panels. Data presented in panels **B** and **C** were collected in a single experiment for each substrate and the TssM and control curves (black and grey) shown in each panel are the same but shown again for clarity. **(D-F)** pH dependence of the dual activities of TssM and the indicated substituted versions. Heatmaps show the area under the curve (AUC) of fluorescence polarization experiments collected using 0.5 µM TssM or its variants, where the higher area correlates to more enzymatic activity. Buffering information can be found in the methods section.

We then subjected these mutant TssM proteins to fluorescence polarization experiments to study their ability to cleave Ub-Lys, Ub-Ser, and Ub-Thr substrates. Single substitutions at either the Asn (NA), His (HA), or Asp (DA) reduced the activity of TssM toward the Ub-Lys substrate to varying degrees, while these versions of TssM retained substantial activity toward the Ub-Ser and Ub-Thr ester-linked substrates at both 0.5 µM and 2 µM enzyme concentration (Figure 2B, Supplemental Figure 1A).

In an attempt to further eliminate TssM Ub esterase activity, we made several double and triple substitutions within the catalytic triad and oxyanion hole. For these substitutions, we focused on the Cys and His positions as the two most important sites for canonical isopeptidase activity. The double mutant CH/AA and the triple mutant NCH/AAA each abolished isopeptidase activity to the same extent as the CA substitution at 0.5 µM and 2 µM enzyme concentration (Figure 2C, Supplemental Figure 1B). Interestingly, only the CH/AA version of TssM showed indications of interaction with the Ub-Lys substrate, providing evidence that Asn286 is important for binding of an isopeptide-linked substrate (Supplemental Figure 1B). The CH/AA double mutant also nearly eliminated activity toward the ester-linked substrates. These results indicate that TssM Cys292 and His426 are independently, but not mutually, dispensable for esterase activity.

### TssM DUB activities exhibit distinct dependencies on pH

Based on the observation of distinct catalytic requirements for isopeptidase and esterase activity, we aimed to study the pH dependence of TssM function. A typical catalytic triad mechanism for cysteine proteases should exhibit a bell-shaped relationship with pH, as the Cys deprotonation is balanced with the proton shuttling of the His residue (Lewis *et al*, 1976, 1981; Polgar, 1974; Storer & Menard, 1994). We observed a clear pH-dependent activity toward the isopeptide Ub-Lys substrate given by the area under the curve (AUC) of fluorescence polarization experiments conducted over pH 4.0-9.2 (Figure 2D). The wildtype TssM enzyme displayed severely impaired isopeptidase activity at pH 4.0 and gradually gained activity as the pH increased, before reaching maximum activity around pH 7.4. Surprisingly, the esterase activity of TssM was not nearly as dependent on pH (Figure 2E, F), as extremely high activity was observed at each pH tested. The individual CA and HA mutants, however, displayed distinct profiles of pH dependence. For the CA mutant, an unusual biphasic pH dependence was observed, wherein the esterase function was slightly less active at pH 8.0 than it was at either pH 7.4 or 8.6. Meanwhile, the HA mutant exhibited a more classical bell-shaped relationship with pH, with a maximal activity at pH 7.2. While the CH/AA double mutant was largely inactive across the pH range, a slight increase in activity was observed at higher pH for only the Ub-Ser substrate. Together, these data support the hypothesis that TssM can utilize differing catalytic mechanisms to support esterase and isopeptidase activities.

### Crystal structures of TssM harbor a water in the catalytic site

The pH dependence and reliance upon the catalytic triad His for Cys-independent esterase activity suggested the possibility of direct water-mediated hydrolysis. To visualize this possibility, we looked to the catalytic site within the apo TssM crystal structure solved previously (Hermanns *et al*., 2024). We observed unmodeled electron density near the catalytic triad in both copies of TssM within the asymmetric unit. After modeling waters into these extra densities, we noted no clashes or other issues following refinement (Figure 3A). When the catalytic Cys292 was aligned in both monomers, we noticed that the sidechain of His426 occupied slightly different positions, indicating flexibility within the active site in this apo state. Similarly to the positional differences in the His side chain, our newly modeled water molecules were 5.6 Å and 7.3 Å away from their respective His426. Conversely, in a crystal structure of Ub-bound TssM (Hermanns *et al*., 2024), a water molecule has already been modeled within the active site of both copies in the asymmetric unit (Figure 3B). In both instances, the water molecule occupies an almost identical position, located 2.7 Å away from His426 and ∼3.4 Å from the Ub C-terminus that would be the site of nucleophilic attack for hydrolysis. Binding of the Ub C-terminal tail into the active site may therefore position the water molecule into hydrogen-bonding distances required for catalysis.

**Figure 3.**
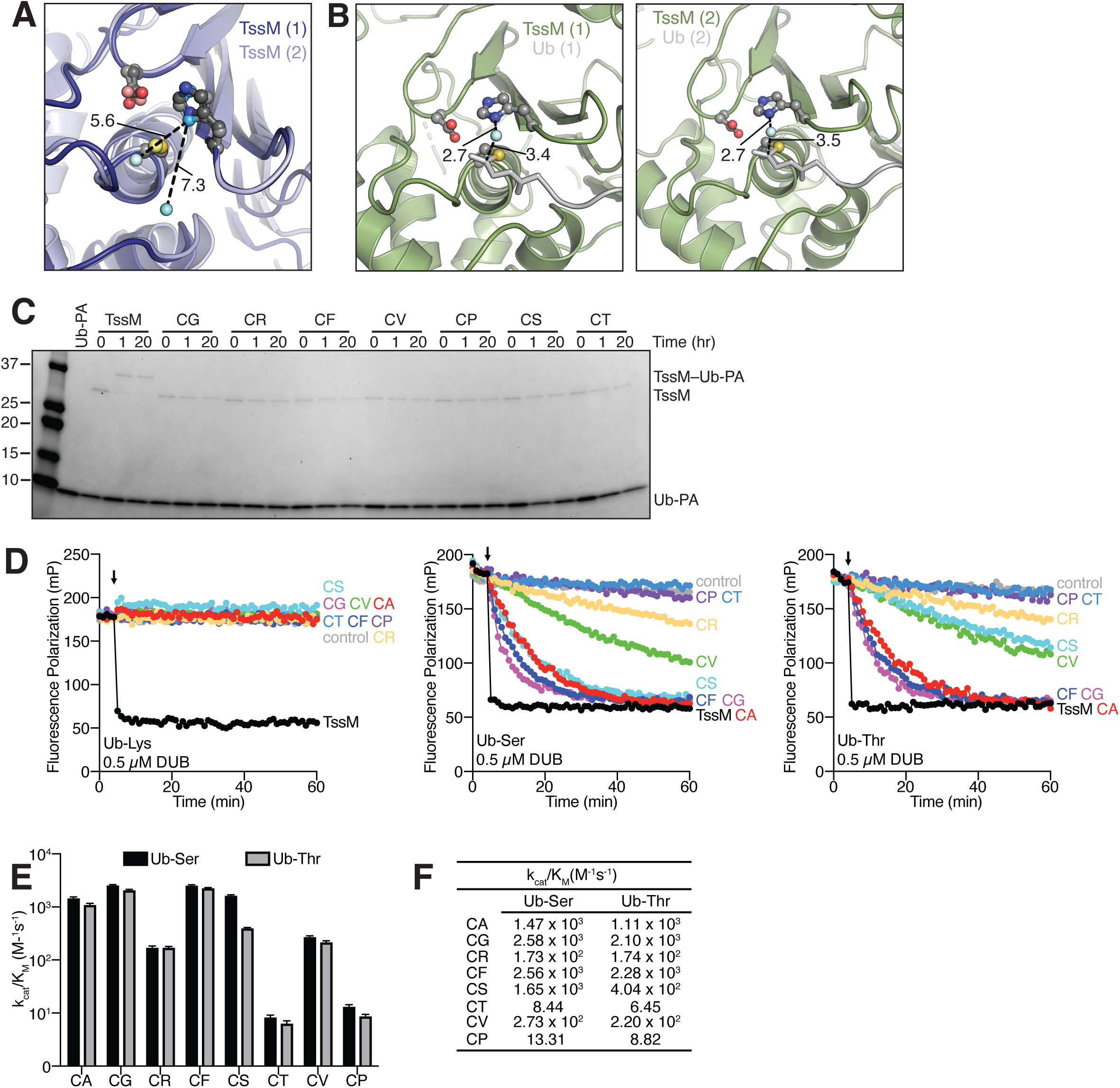
The catalytic Cys is dispensable for the esterase activity of TssM. **(A)** Structure of apo TssM with monomers in the asymmetric unit aligned at Cys292 (PDB: 8PZ3). Previously unmodeled electron density was modeled with water and refined. Distances between the catalytic His and the water (cyan spheres) are shown by the black dashed lines. Monomers are colored such that the darker colors correlate to one monomer and the lighter colors show the other monomer. **(B)** Structures showing each TssM-Ub in the asymmetric unit (PDB: 8Q00). Water molecules are indicated by cyan spheres, and distances between the catalytic His, the water, and the C-terminus of Ub are shown by dashed lines. **(C)** Ub-PA reactions with the indicated catalytic Cys versions of TssM. **(D)** Fluorescence polarization experiments for Cys versions of TssM at 0.5 µM enzyme concentration toward the Ub-Lys, Ub-Ser, and Ub-Thr substrates. In each panel, DUB addition is marked by a black arrow. **(E, F)** Mean catalytic efficiencies (k_cat/_K_M_) for each Cys version of TssM. Values for the CA version of TssM are shown again from Figure 1 for ease of comparison.

### Several Cys variants of TssM retain esterase activity

While trying to create an entirely inactive version of TssM, we made several additional point substitutions at Cys292. In addition to the Ser and Arg substitutions used in prior DUB studies (Morrow *et al*., 2018), we further explored elimination of the side chain with a glycine (Gly, G) substitution, as well as increasing bulk with threonine (Thr, T), valine (Val, V), proline (Pro, P), and phenylalanine (Phe, F) substitutions. As expected, the Ub-PA probe was unable to covalently modify these Cys variants of TssM (Figure 3C). Several Cys substitutions, including CR, CF, CV, and CP, resulted in a slight destabilization of the TssM fold, as evidenced by up to a ∼7-degree reduction in melting temperature (Supplemental Figure 2A). Meanwhile, the CS version of TssM exhibited slightly higher stability, evidenced by a ∼4-degree increase in melting temperature (Supplemental Figure 2A). Finally, TssM versions including CA, CG, and CT displayed no significant change in melting temperature, demonstrating their limited impact on the overall fold (Supplemental Figure 2A). None of the tested Cys variants displayed any activity toward the isopeptide-linked Ub-Lys substrate (Figure 3D, Supplemental Figure 2B). Against the ester-linked Ub-Ser and Ub-Thr substrates, however, this panel of substitutions displayed a range of diminished activities compared to wildtype TssM. The CG and CF versions of TssM retained similar levels of activity as the CA mutant toward both ester-linked substrates. The CS, CV, and CR versions exhibited reduced activity compared to the CA mutant, though the decrease observed in the CS mutant was specific to the Ub-Thr substrate. Lastly, only the CT and CP substitutions in TssM could virtually eliminate esterase activity (Figure 3D, Supplemental Figure 2B).

### TssM Cys substitutions reduce catalytic efficiency toward ubiquitin esters

We previously showed that the catalytic efficiency of wildtype TssM toward Ub-Ser and Ub-Thr fluorescence polarization substrates was approaching 10^7^ M^-1^s^-1^ (Szczesna *et al*., 2024). We determined the catalytic efficiency of each individual TssM Cys variant, and all versions showed drastic reduction in the k_cat_/K_M_, (Figure 3E, F). Like the CA version of TssM shown earlier, the CG, CF, and CS versions of TssM reduced catalytic efficiency by ∼10^4^, while CR and CV versions reduced ∼10^5^. The CT and CP versions of TssM displayed the strongest reduction in activity, showed by their k_cat_/K_M_ measuring only ∼10 M^-1^s^-1^, marking a nearly 10^6^ reduction in the efficiency of TssM to cleave either Ub-Ser or Ub-Thr substrates. Thus, select Cys substitutions are far superior to others at eliminating TssM Ub esterase activity.

### The CT substitution eliminates isopeptidase and esterase activity while maintaining the stability of TssM

Our fluorescence polarization data indicate that substitution of the catalytic Cys to a Thr significantly reduces all enzymatic activity of TssM (Figure 3E, F). AlphaFold3 models the USP domain of the TssM CT mutant with overall very high confidence, with some areas of high confidence (Figure 4A, Supplemental Figure 2C). Modeling the CT version of TssM with a Ub molecule using AlphaFold3 increased the confidence throughout the USP domain and maintained the orientation of the Thr292 sidechain compared to the apo model (Figure 4B, Supplemental Figure 2D, E) The structure around the active site was modeled with very high confidence in both models, but AlphaFold3 was somewhat less confident about the side chain position for Thr292 and His426 (Supplemental Figure 2C, D). Closer inspection of the active site revealed that all resulting models positioned the sidechain hydroxyl of the substituted Thr nearly within hydrogen-bonding distance of the His426 sidechain in both the apo and Ub-bound models (< 4.0 Å) (Figure 4A, B). Thus, the CT version of TssM is predicted to position the active site similarly to the wildtype version of the enzyme.

**Figure 4.**
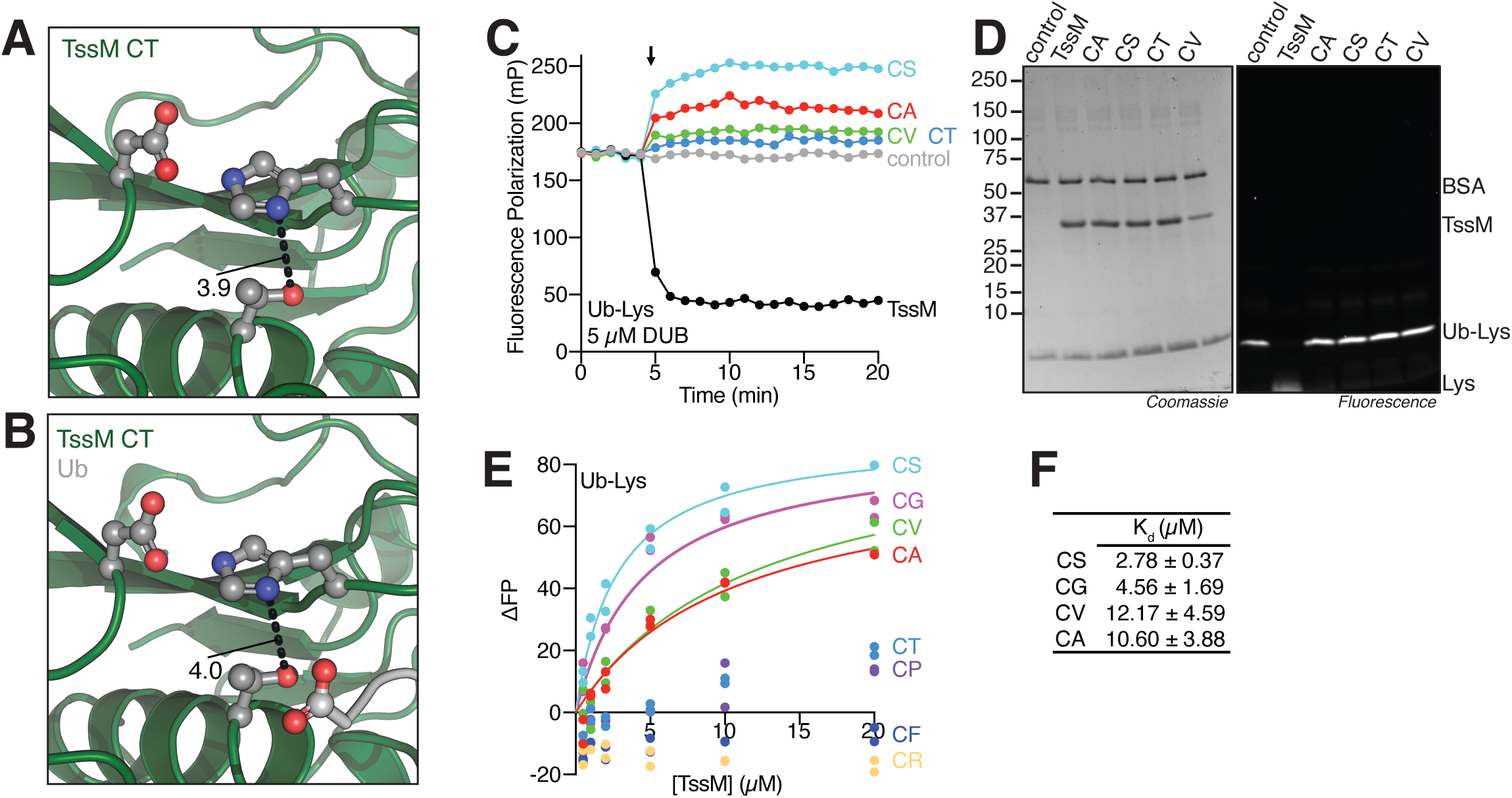
Cys versions of TssM disrupt the formation of the first tetrahedral intermediate required for isopeptidase activity. AlphaFold3 modeling showing the active site of the **(A)** CT version of TssM and **(B)** TssM CT with Ub bound. **(C)** Fluorescence polarization experiment showing variants of TssM noncovalently bind the Ub-Lys substrate at 5 µM. **(D)** Samples from (C) were removed from the plate after the 20-minute experiment and species were resolved using SDS-PAGE. Shown are Coomassie staining and fluorescence scans of the gel. **(E)** Fluorescence polarization binding experiment for Ub-Lys with the indicated versions of TssM. Curves represent a one-site total binding fit, and fits for datasets without curves could not be determined. **(F)** K_d_ values and associated standard error of the mean (SEM) corresponding to curves that could be fit in (E).

### Cys-substituted versions of TssM are deficient in forming the tetrahedral intermediate required for isopeptidase activity

Canonical Cys-based isopeptidase activity requires the formation of a tetrahedral intermediate that is stabilized by an oxyanion hole (Mevissen & Komander, 2017). Following resolution of the tetrahedral intermediate and release of substrate, the DUB catalytic Cys remains covalently modified with the Ub in an acyl-enzyme intermediate. Subsequently, water-mediated hydrolysis liberates the covalently-bound Ub from the Cys to regenerate an active enzyme. To determine which step in the canonical isopeptidase mechanism Cys-substituted versions of TssM are deficient, we incubated a high enzyme concentration of TssM wildtype or the CA, CS, CT, or CV mutants with the Ub-Lys substrate and monitored fluorescence polarization. As shown for lower enzyme concentrations, only wildtype TssM could cleave the Ub-Lys substrate (Figure 4C). Addition of each Cys-substituted version of TssM instead caused the fluorescence polarization signal to increase, indicative of a larger complex forming in solution with the Ub-Lys substrate. To determine whether this complex was noncovalent or indicative of some unresolved, Cys-independent DUB∼Ub covalent intermediate, we removed samples from the fluorescence polarization experiment at the end of the time course (20 minutes) and evaluated them by SDS-PAGE (Figure 4D). We observed no higher molecular weight fluorescence that would indicate a stable covalent intermediate between with TssM or its Cys variants and the Ub-Lys substrate. The wildtype TssM reaction was the only treatment that successfully liberated the fluorophore from the C-terminus of Ub, a direct result of the enzyme’s isopeptidase activity. Accordingly, this result indicates that 1) the inability of TssM Cys variants to cleave a canonical isopeptide-linked substrate results from their lack of a nucleophile required to form a productive tetrahedral intermediate in the active site; and 2) TssM esterase activity does not require the formation of the canonical tetrahedral intermediate to promote catalysis.

### Large, bulky substitutions at position 292 exclude ubiquitin from the active site

With the knowledge that all Cys versions have lost their nucleophilic capacity, we sought to better understand why some substitutions better eliminate esterase function than others. To do so, we took advantage of the lack of isopeptidase capacity to measure binding of the Ub-Lys substrate into the substituted TssM active sites by fluorescence polarization (Figure 4E). Due to the proximity of the fluorophore to the Ub C-terminus, this measure should directly report on even subtle changes in binding at the active site. Small substitutions such as CS, CG, CA, and CV, which all retain some Ub esterase function, permitted the binding of Ub-Lys with K_d_ values in the low micromolar range (Figure 4F). Both CT and CP versions showed some evidence of binding at higher enzyme concentrations but were unable to be fit for K_d_ determination. Finally, we observed a net negative change in the fluorescence polarization signal for the CR and CF versions of TssM, compared to the control signal of just Ub-Lys in solution. This unusual result is consistent with an even higher degree of flexibility imparted by exclusion from the TssM active site. Therefore, small substitutions within the active site permit the binding of a ubiquitinated substrate, whereas large, bulky substitutions exclude the C-terminus of Ub from the active site.

### Some reportedly inactive human DUBs exhibit ubiquitin esterase activity

To extend our analysis of Ub esterase catalytic requirements to human DUBs, we evaluated USP21 and JOSD1, two DUBs with known dual or selective esterase activity, respectively (De Cesare *et al*., 2021). Additionally, we screened several human DUBs with no reported activity including OTUD6B, USP50, ALG13, and the pseudo-DUB OTULINL. These DUBs had previously only been evaluated for isopeptidase activity, with the exception of OTUD6B, which is reportedly inactive also against a Ub-Thr substrate (Ceccarelli *et al*., 2019; De Cesare *et al*., 2021; Mevissen *et al*., 2013; Ritorto *et al*., 2014). Evaluating the selected DUBs against each substrate at 0.5 µM enzyme concentration shows that USP21 is the only dual isopeptidase/esterase DUB in our panel, while JOSD1 displays preferential esterase activity toward the Ub-Ser and Ub-Thr substrates (Figure 5A). Surprisingly, OTUD6B displayed marked activity exclusively toward the Ub-Ser substrate, in a concentration-dependent manner (Figure 5A, Supplemental Figure 3A). At 2 µM enzyme concentration, OTUD6B displayed Ub-Thr binding, evidenced by the increase in fluorescence polarization after OTUD6B addition (Supplemental Figure 3A). We also examined a version of OTUD6B expressed and purified from insect cells (iOTUD6B) at both 0.5 µM and 2 µM enzyme concentration and observed an even higher level of activity toward Ub-Ser, along with low-level isopeptidase activity at 0.5 µM that became more apparent at 2 µM (Supplemental Figure 3B, C). ALG13 and USP50 were inactive at both 0.5 µM and 2 µM enzyme concentration toward all substrates tested. OTULINL displayed weak activity toward the Ub-Ser substrate at 2 µM enzyme concentration, highlighting the first evidence of Ub esterase activity in this pseudo-DUB that lacks a conserved catalytic Cys (Supplemental Figure 3A). In light of our observations that Ub isopeptidase and esterase activities have distinct catalytic requirements, these results demonstrate the importance of screening DUBs against all possible substrates, as activity for both OTUD6B and OTULINL had not been observed previously against canonical isopeptide-linked substrates.

**Figure 5.**
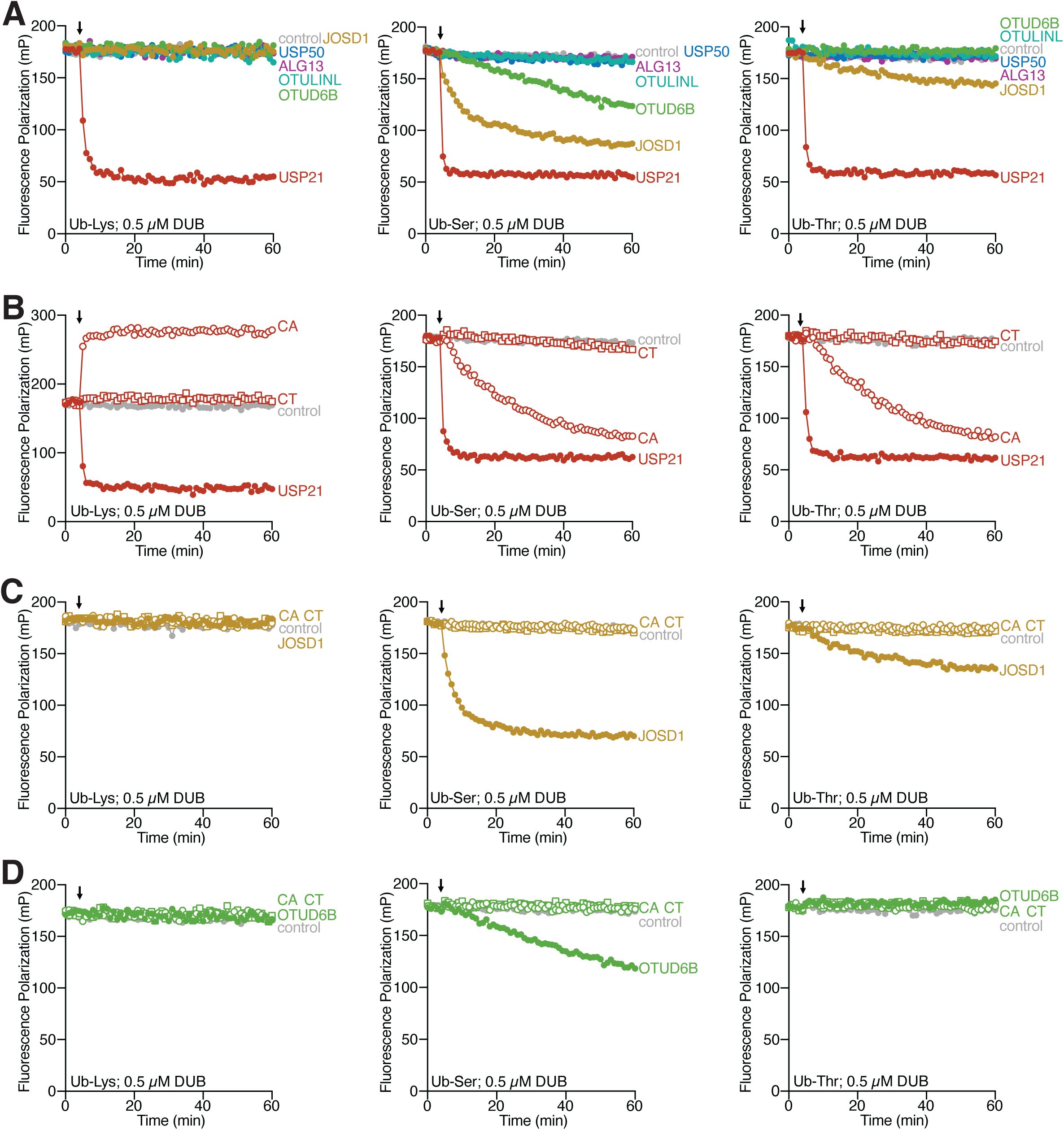
Some reportedly inactive human DUBs exhibit ubiquitin esterase activity. **A)** Fluorescence polarization experiments showing the activity of the indicated DUBs at 0.5 µM enzyme concentration toward the Ub-Lys, Ub-Ser, and Ub-Thr substrates. The impacts of substitution at the catalytic Cys (CA or CT) for **(B)** USP21, **(C)** JOSD1, and **(D)** OTUD6B at 0.5 µM enzyme concentration are shown. For these panels, the wildtype version of the protein is indicated by solid circles, the CA version is indicated by open circles, and the CT version is indicated by the open squares. In each panel, DUB addition is marked by a black arrow.

### CT substitutions of human DUBs also eliminate esterase activity

Our observation that the CA variant of TssM retains Ub esterase activity prompted us to evaluate whether the CA versions of other DUBs also retained activity. For the dual isopeptidase/esterase activity of USP21, the CA version of the protein was indeed still active against the Ub-Ser and Ub-Thr substrates but failed to cleave the Ub-Lys substrate (Figure 5B, open circles). Similarly to TssM, the CT version of USP21 abolished both types of DUB activity at 0.5 µM enzyme concentration (Figure 5B, open squares). At higher enzyme concentration, however, even the CT version of USP21 demonstrated some low-level esterase activity (Supplemental Figure 3D). Based on these findings with USP21 and the bacterial USP-type DUB TssM, the CA substitution of other dual isopeptidase/esterase DUBs within the USP family may be insufficient to completely abolish activity.

We also examined the CA and CT versions of the ubiquitin esterases JOSD1 and OTUD6B (Figure 5C, D; Supplemental Figure 3E, F). For these enzymes, the CA substitution was sufficient to eliminate the catalytic activity, and there was no added benefit to utilizing a CT substitution. Together, these results indicate that while a CA substitution may sufficiently eliminate esterase activity in some DUBs, a CT substitution could be more broadly effective.

## Discussion

In this study we show that for the dual isopeptidase/esterase DUB TssM, a standard Cys-to-Ala substitution at the catalytic site is insufficient to abolish activity. We show that the CA version of TssM only eliminates its isopeptidase activity, but the enzyme retains considerable activity toward ester-linked substrates. The catalytic efficiency of the CA version of TssM toward both Ub-Ser and Ub-Thr substrates is on the order of 10^3^ M^-1^s^-1^ (Figure 3E, F), which is comparable to several other isopeptidase DUBs including Trabid (K29 or K33 Ub dimers; (Geurink *et al*, 2016)) USP30 (Ub-Lys; (Gersch *et al*., 2017)), and USP1, USP21, USP30 (Ub-AMC; (Faesen *et al*, 2011)). Further, the catalytic efficiency of JOSD1 toward ester-linked Ub substrates was shown to be between 10^3^ and 10^4^ (De Cesare *et al*., 2021; Szczesna *et al*., 2024), and the catalytic efficiencies for several metalloprotease DUBs, which utilize a similar Cys-independent mechanism, are all on the order of 10^4^ (K63 Ub dimer; (Sato *et al*, 2008; Shrestha *et al*, 2014; Zeqiraj *et al*, 2015)). These comparisons highlight the significant level of residual esterase activity observed in the TssM CA version.

Our analyses have demonstrated distinct catalytic requirements for DUB functions and provided a basis for proposing the mechanism of Cys-independent esterase activity. Namely, in the absence of a nucleophilic active site Cys, a water molecule is activated by the nearby His to permit direct hydrolysis of an ester-linked Ub species. This explains why the catalytic Cys and His residues are independently but not mutually dispensable for esterase activity (Figure 2B, C) and is consistent with the observed pH-dependence (Figure 2D-F). It is also supported by the inability of TssM Cys mutants to form a covalent intermediate with the Ub-Lys substrate (Figure 4C, D). Further, several DUB structures (including those for TssM, Figure 3A, B) show at least one water molecule near the catalytic His side chain (Hermanns *et al*., 2024; Keijzer *et al*, 2024) demonstrating that water penetrates the DUB active site as part of resolving the acyl-enzyme intermediate during the isopeptidase catalytic cycle. We propose that, in the absence of a catalytic Cys residue, this same water molecule can be activated by the nearby His for direct hydrolysis of ester-linked Ub substrates (Figure 6).

**Figure 6.**
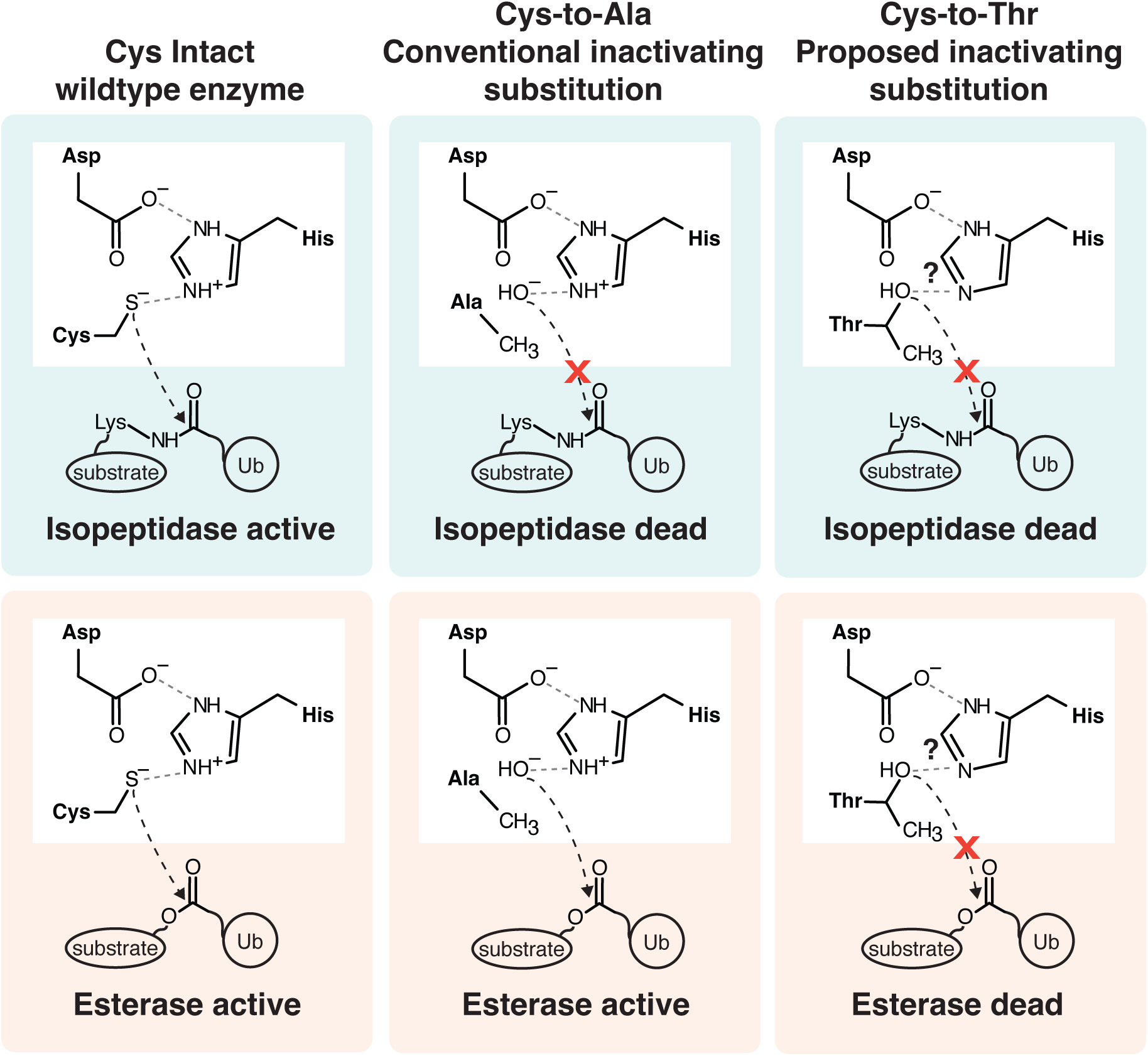
Proposed mechanisms for isopeptidase and esterase inactivation in dual activity DUBs. Shown are schematics representing the wildtype version of a DUB capable of both isopeptidase and esterase activities, the Cys-to-Ala version commonly used as an inactivating substitution that fails to eliminate esterase activity, and the Cys-to-Thr version capable of eliminating both activities.

This underlying capacity for direct water-mediated hydrolysis of Ub esters poses a new challenge for mutational studies. In addition to the complications raised herein with the common Ala substitution, prior work studying canonical isopeptide-linked Ub modifications has presented the risk that CA variants can behave in a dominant-negative fashion by retaining binding capacity (Damgaard *et al*, 2016; Morrow *et al*., 2018). In some cases, this has been addressed by incorporating a bulky Arg substitution instead, in an attempt to exclude the Ub C-terminus from the active site. In our studies with TssM, we observe that bulky substitutions with either Arg or Phe indeed appear to perturb binding within the active site, but do not fully abrogate Ub esterase activity. A Pro substitution eliminated both binding and activity, but likely through local structural destabilization, as indicated by a decreased melting temperature. A Thr substitution appeared to retain folding, eliminate binding, and largely abolish esterase activity. Based on these observations, we postulate that a Thr substitution serves three roles: 1) elimination of nucleophilic capacity, 2) addition of steric bulk, and 3) addition of hydrogen-bonding capability to disrupt contacts to the catalytic His or other nearby hydrogen-bond acceptors that could otherwise stabilize and activate a water molecule. A Thr substitution in USP21 was also much more potent in abolishing its esterase activity, suggesting that at least among USP-type DUBs, this may be a preferred option for future mutational studies.

Our work presents two important considerations as the field of non-canonical ubiquitination continues to expand. First, use of the conventional inactivating mutation (Cys-to-Ala) may not be the best option for DUB functional studies due to its inability to inactivate the esterase activity of dual isopeptidase/esterase enzymes such as TssM and USP21. Second, with the newfound variety of ubiquitination substrates, including non-proteinaceous molecules (Jochem *et al*., 2026; Kelsall *et al*., 2022; Otten *et al*., 2021; Wang *et al*., 2025; Yip *et al*., 2026; Yoshida *et al*., 2024), it becomes increasingly more important to evaluate the activity of a DUB against all available substrates. OTULINL, which was previously classified as a pseudo-DUB because it lacked the catalytic requirements established for isopeptidase function (Ceccarelli *et al*., 2019), harbors activity toward the ester-linked Ub-Ser substrate (Supplemental Figure 3A). Likewise, our analysis also identified a cryptic Ub esterase activity in the OTU family DUB, OTUD6B, which is largely specific for Ub-Ser. In both OTULINL and OTUD6B, the identification of Ub esterase functions could provide a molecular explanation for cancer- and neurodevelopment-associated missense variants mapping to the OTU domain (The Cancer Genome Atlas (TCGA), (Cingoz *et al*, 2022; Santiago-Sim *et al*, 2017; Straniero *et al*, 2018)). In all, this work offers yet another reminder that as our appreciation for the complexity of ubiquitin biology expands, we must continuously evaluate available tools and underlying assumptions of our approaches.

## Materials and Methods

### Plasmids and cloning

The DNA sequence for *Burkholderia pseudomallei* TssM encoding the Big5 and USP domains (residues 192-474) was generated as described previously (Szczesna *et al*., 2024). pOPINK-OTUD6B was purchased from Addgene (#61418). pET28b-JOSD1 was a gift from S. Buhrlage (Dana Farber). DNA sequences for human OTULINL, ALG13, and USP50 were synthesized by Twist Biosciences and subsequently cloned into the pOPINB expression vector. The full-length sequences for OTUD6B, JOSD1, OTULINL and USP50 and the ALG13 OTU domain (216-353) were used. All point substitutions were generated by site-directed mutagenesis and confirmed by DNA sequencing. Sequences for several of the TssM mutants (for example, CA and CV) were obtained after multiple passages of plasmids to validate the purity.

### Protein expression and purification

All plasmids were transformed and expressed from Rosetta *E. coli* cells after a fresh transformation. A single colony was selected and grown in a starter culture overnight at 37 °C with the appropriate antibiotics supplemented in LB media. The starter cultures were used to inoculate larger growths containing antibiotics, and the bacteria were grown at 37 °C until the OD_600_ reached ∼0.8. The temperature was reduced to 18 °C and protein expression was induced with the addition of 0.5 mM IPTG overnight. Cell pellets were harvested and resuspended in 25 mM Tris pH 7.4, 200 mM NaCl prior to lysis by sonication. Lysates were clarified by centrifugation at 45000g for 1 hour. Clarified lysates were passed through Ni-NTA affinity resin and subsequently washed with 25 mM Tris pH 7.4, 200 mM NaCl at two steps of imidazole (10 mM and 35 mM). Bound proteins were eluted using 25 mM Tris pH 7.4, 200 mM NaCl, 250 mM imidazole. During dialysis back to 25 mM Tris pH 7.4, 200 mM NaCl, TEV protease was used to remove the tag from all TssM versions, while 3C protease was used to remove the His-GST tag from OTUD6B and the His tags from OTULINL, USP50, and ALG13. JOSD1 retains an N-terminal His tag, and USP21 retains an N-terminal His-SUMO tag. For DUBs that had affinity tags removed, the cleaved protein was passed back through Ni-NTA resin and the flowthrough was collected and concentrated. The final purification for all enzymes regardless of tag removal was by size exclusion chromatography (HiLoad 16/600 Superdex75 for all versions of OTULINL, USP50, JOSD1, and OTUD6B; or HiLoad 16/600 Superdex200 for all versions of TssM and USP21) in 25 mM Tris pH 7.4, 150 mM NaCl, 5 mM DTT. The insect preparation of OTUD6B was achieved by expression in SF9 cells followed by purification by Ni-NTA affinity chromatography. In the case of wildtype iOTUD6B, an additional purification step was performed using size exclusion chromatography using a HiLoad 16/600 Superdex75 in 50 mM Tris pH 8, 100 mM NaCl, 1 mM DTT, and 1 mM EDTA. Proteins were then quantified by absorbance at 280 nm, aliquoted, and flash frozen for storage at -80 °C.

### Ub-Propargylamide (Ub-PA) reaction

Reactions were conducted using 0.5 µM TssM, 20 µM Ub-PA, and 5 mM DTT made up in 50 mM Hepes pH 7.4, 100 mM NaCl. Tubes were incubated at 22 °C for the duration of the experiment, and timepoints were quenched in 3x SDS-PAGE sample buffer. Protein species were resolved using a 4-20% Criterion TGX Precast Midi protein gel (BioRad) and visualized by Coomassie blue staining. For the experiment shown in Figure 1D, 5 µM TssM was used.

### Fluorescence polarization (FP) experiments

TAMRA-K(Ub)G (Ub-Lys), Rho-S(Ub)G (Ub-Ser), and Rho-T(Ub)G (Ub-Thr) FP substrates were prepared as described previously (Geurink *et al*, 2012; Szczesna *et al*., 2024). FP was monitored using a BMG Labtech CLARIOstar microplate reader. Reaction volumes were 20 µL and data were collected in a black, low-protein binding 384-well plate at 22 °C, blanked against fresh FP buffer containing 50 mM HEPES (pH 7.4), 100 mM NaCl, 0.1 mg/mL BSA, 5 mM β-mercaptoethanol. All experiments except the pH titration data shown in Figure 2D were collected in this FP buffer. For the pH titration, the buffering agents were NaOAc (pH 4.0, 4.6, 5.2), BisTris (pH 5.8, 6.4, 7.2), HEPES (pH 7.4, 8.0), and CHES (pH 8.6, 9.2). For the ester-linked substrates conjugated to a rhodamine110 fluorophore, 482-16 nm and 530-40 nm optic filters were used for excitation and emission, respectively. For the isopeptide-linked Ub-Lys substrate, the Tamra fluorophore was excited at 540-20 nm and emission was monitored at 590-20 nm. In each reaction, the fluorescent substrate is present at 50 nM. The concentration of DUB used in each experiment ranged from 0.5 µM to 5 µM except for the kinetic experiments, where concentration panels were optimized to derive k_cat_/K_M_ values as described below. All reactions were performed in duplicate, and data were processed following blank subtraction.

For kinetic experiments, a range of 6 different concentrations were selected for each protein based on its level of activity. The highest concentration range used was for TssM CT (5 to 50 µM), and the lowest concentration range was for TssM CS (0.2 to 2 µM). The FP curves obtained were then analyzed by non-linear regression using a one-phase decay fit from GraphPad Prism 10. Rate constants (k) were determined for each concentration by constraining k to be shared between all replicates (n = 2). Rate constants and the associated standard error of the mean (SEM) were extracted and graphed against the concentration of DUB. Repeating this processing over the range of concentrations yielded linear fits that corresponded to the k_cat_/K_M_ (in M^-1^s^-1^; catalytic efficiency).

For binding experiments to derive the K_d_, versions of TssM were incubated with 100 nM Ub-Lys at concentrations ranging from 0.5 µM to 20 µM. Five reads were collected and averaged to attain an FP signal for each DUB at each concentration. Change in FP signal (ΔFP) was determined by subtracting the signal for the negative control well (substrate only) from each measurement with TssM. Data were then graphed as a function of concentration and fit to a one-phase total binding curve using GraphPad Prism 10.

### Thermal stability assay

A thermal stability assay for TssM variants was conducted in MicroAmp Fast 96-Well Reaction Plates (Applied Biosystems) with SYPRO Orange Protein Gel Stain (MilliporeSigma) using a QuantStudio 3 Real-Time PCR system (Applied Biosystems). Assays were performed in 20 µL volumes containing 10 µM TssM variants and 20× SYPRO dye (diluted from a 5000× stock) in 50 mM HEPES pH 7.4, 100 mM NaCl. The protocol increased the temperature from 22 to 99 °C on a gradient of 0.1 °C/s, and fluorescence was monitored using an excitation wavelength of 580 ± 10 nm and an emission wavelength of 623 ± 14 nm.

### Structural analysis

Structural modeling of TssM CT was performed using the AlphaFold3 web interface (Abramson *et al*, 2024). AlphaFold3 PAE plots were visualized using PAE Viewer (Elfmann & Stulke, 2023). Structural analysis and figure generation were performed using PyMol (The PyMOL Molecular Graphics System, Version 3.0 Schrödinger, LLC).

## Data Availability

This study includes no data deposited in external repositories. Reagents will be made available upon request to the corresponding author.

## Acknowledgements

We wish to thank Dr. Christopher Lima as well as members of the Pruneda Lab for their helpful suggestions to strengthen our experimental design and data analysis. We thank Dr. Achim Werner for OTUD6B reagents. This work was supported by the National Institute of General Medical Sciences (R35GM142486 to JNP).

---

**Supplemental Figure 1.**
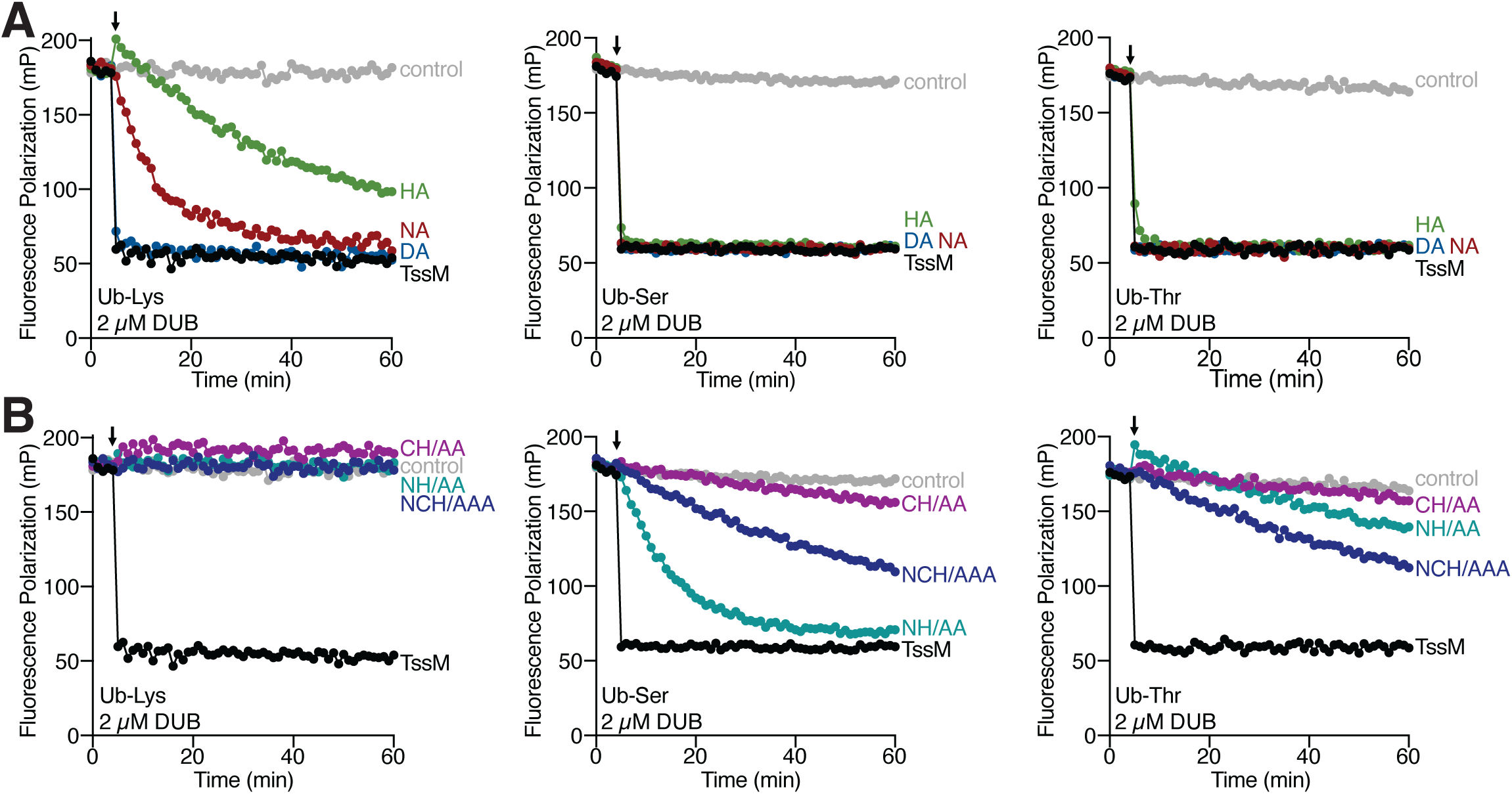
C**y**s292 **or His426, individually, are sufficient to promote esterase activity in TssM.** Fluorescence polarization experiments showing the activity of **(A)** single catalytic substitutions, and **(B)** double and triple catalytic substitutions within TssM towards Ub-Lys, Ub-Ser, and Ub-Thr substrates at 2 µM enzyme concentration. In each panel, DUB addition is marked by the black arrow. All data presented were collected in a single experiment per substrate, and the data shown between panels for wildtype TssM (black) and the control (grey) are the same for each substrate but shown twice for clarity.

**Supplemental Figure 2.**
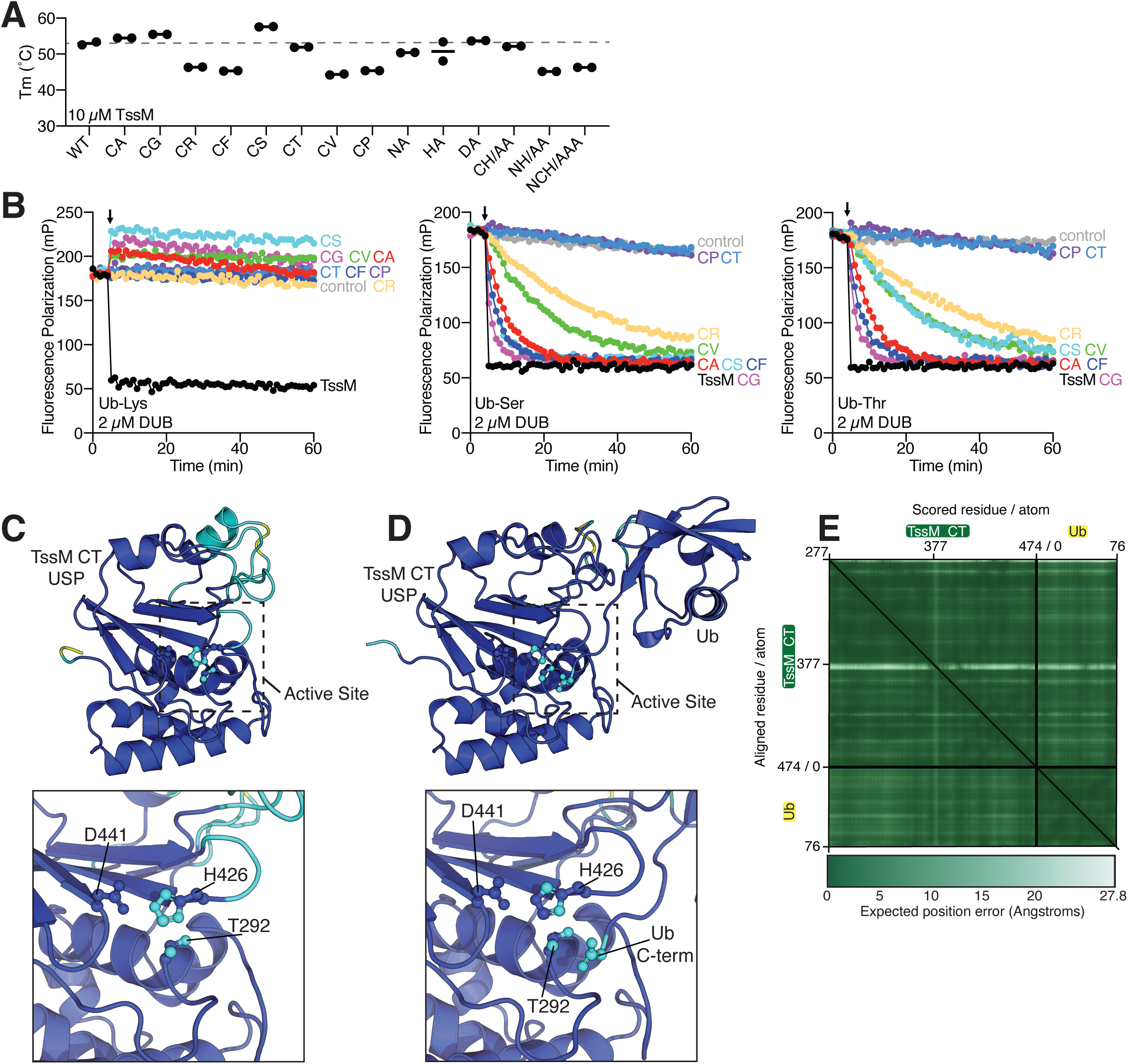
The catalytic Cys is dispensable for the esterase activity of TssM. **(A)** Thermal stability assay showing the melting temperature of TssM and its substituted versions used in this study. **(B)** Activity of the catalytic Cys versions of TssM at 2 µM enzyme concentration toward Ub-Lys, Ub-Ser, and Ub-Thr substrates. DUB addition is marked by a black arrow. AlphaFold3 model of **(C)** TssM CT and **(D)** TssM CT with Ub, including close-up views of the active site colored by AlphaFold3 confidence. **(E)** PAE plot from AlphaFold3 of the noncovalent TssM CT–Ub model. Darker green indicates higher confidence in the model due to a lower expected position error. For all structure panels, AlphaFold3 coloring is defined by confidence where blue represents pLDDT > 90, cyan 70 > pLDDT > 90, yellow 50 > pLDDT > 70, and orange pLDDT < 50.

**Supplemental Figure 3.**
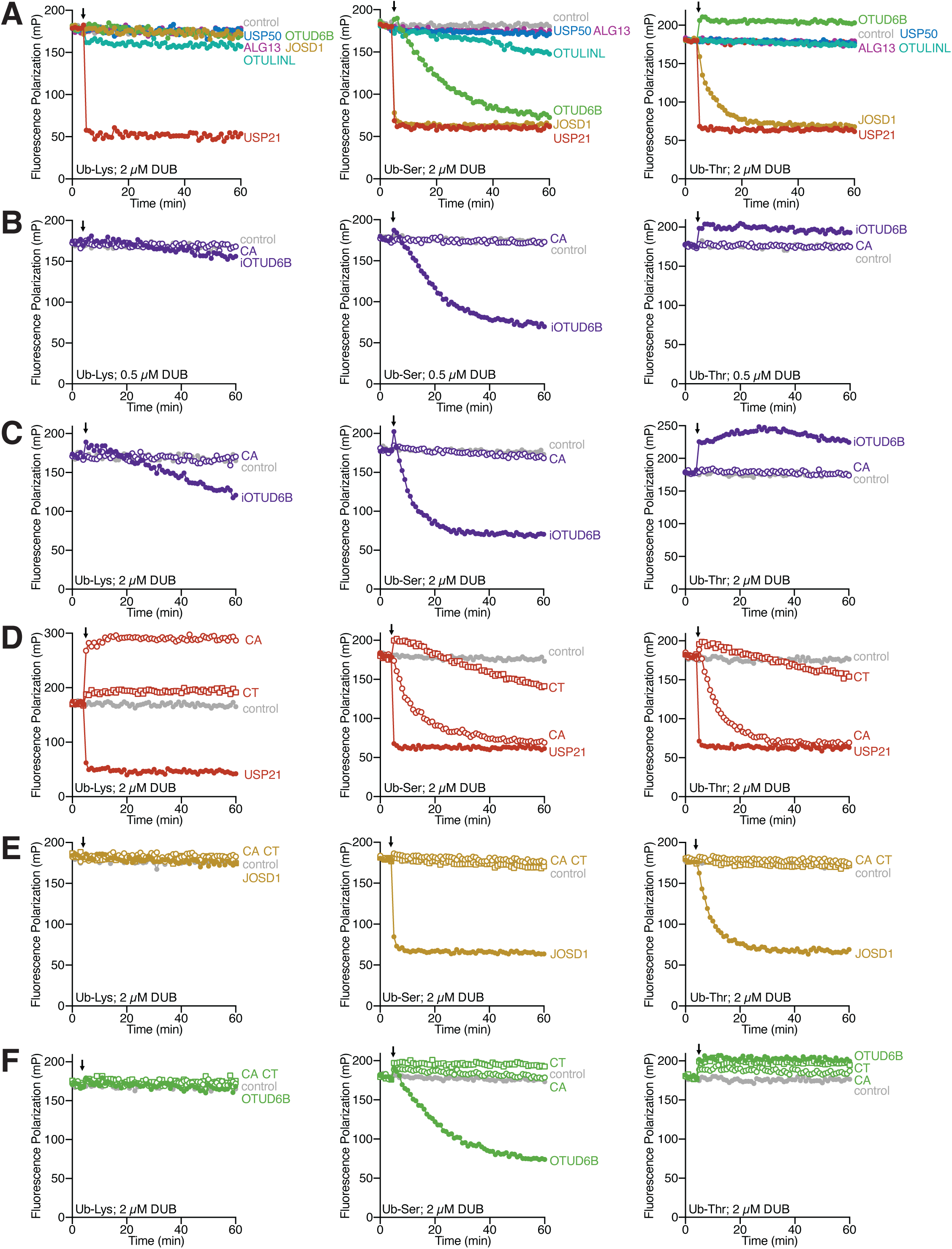
Some reportedly inactive human DUBs exhibit ubiquitin esterase activity. **(A)** Fluorescence polarization experiments showing the activity of the indicated DUBs at 2 µM enzyme concentration toward the Ub-Lys, Ub-Ser, and Ub-Thr substrates. An SF9-expressed version of OTUD6B was evaluated against each fluorescence polarization substrate at **(B)** 0.5 µM enzyme concentration, and **(C)** 2 µM enzyme concentration. The impacts of substitution at the catalytic Cys (CA or CT) for **(D)** USP21, **(E)** JOSD1, and **(F)** OTUD6B at 2 µM enzyme concentration are shown. For these panels, the wildtype version of the protein is indicated by solid circles, the CA version is indicated by open circles, and the CT version is indicated by the open squares. DUB addition is always marked by a black arrow.

